# *In vivo* multi-site electrophysiology enabled by flexible optrodes towards bi-directional spinal cord interrogation

**DOI:** 10.1101/2023.09.20.558602

**Authors:** Pietro Metuh, Marcello Meneghetti, Rune W. Berg, Christos Markos

**Affiliations:** Department of Electrical and Photonics Engineering, Technical University of Denmark, Bygning 343, Ørsteds Plads, DK-2800 Kgs. Lyngby, Denmark; Department of Neuroscience, University of Copenhagen, Blegdamsvej 3B, DK-2200 Kbh N, Denmark

## Abstract

Optical neural interfaces combining optogenetics and electrophysiology have been demonstrated as powerful tools for distinguishing the causal roles of neural circuits in the nervous system. Functional optrodes for multipoint stimulation and recording have already been demonstrated in the brain. However, soft and flexible multimodal optrodes for the purpose of probing the spinal cord have remained undeveloped. Here, we present the design and fabrication of a novel optrode for multi-site optical stimulation and electrical recording in the spinal cord by combining optical fiber drawing of polymer material, laser micromachining, and integration of tungsten microelectrodes in a monolithic fiber-based structure. The results from space-resolved scattering measurements, electrochemical impedance spectroscopy, and an acute *in vivo* electrophysiology experiment in an anesthetized rodent suggest this probe as a potential novel interface, which can serve as a part of therapeutic strategies against neurological conditions and injury in the spinal cord.

## 1 Introduction

Multimodal soft and flexible neural probes have been demonstrated as a powerful tool for distinguishing the causal roles of neural regions in the brain by simultaneously stimulating selected regions and recording their neural response [1, 2]. The ability to combine advanced functional materials in a fiber form has enabled the development of novel photonic devices [3–5] as well as neural interfaces integrating several functionalities such as stimulation, recording, drug delivery, sensing and imaging [6– 11]. In particular, the combination of optical stimulation with optogenetics, which exploits light-sensitive ionic membrane channel proteins (opsins), and electrical recording with electrophysiology, which is used to probe and record the local electrical neural response, constitutes an important tool towards the investigation of brain activity [12]. Moreover, optogenetics and electrophysiology are being developed for treating neurodegenerative diseases, such as seizure control [2] and relief in Parkinson’s [13] and Alzheimer’s disease [14]. In particular, the ability to create multiple stimulation and recording sites can help to link the neural activity of small groups of neurons to specific roles in the central nervous system [15]. Although multi-site optogenetic stimulation or electrophysiology in the brain has already been demonstrated [15–18], there is, to our knowledge, no similar neural interface targeting another fundamental component of the central nervous system, the spinal cord. The spinal cord and the peripheral nervous system flex with the body during movement [19]; therefore, the flexibility of the probe is crucial when targeting these regions, which are rich with excitable cells for therapeutic stimulation against conditions such as injuries or epilepsy [20]. This work presents the implementation of multi-site optical stimulation and electrophysiology on a flexible, tissue-compatible substrate made of polycarbonate (PC) that can accommodate the spinal cord shape thanks to its rectangular cross-section. The optrode design consists of a microstructured PC waveguide to deliver light in the tissue, with embedded tungsten (W) microwires that act as electrodes to record the electrical neuronal response. The PC waveguide has a rectangular cross-section and four aligned microchannels, inside which tungsten microwires of 50 µm in diameter are inserted for electrical recordings. The electrodes are exposed by laser micromachining one hole per channel on the surface of the waveguide. Simultaneously, the micro-holes introduce scattering points for the propagating light so that the optical scattering is enhanced and the tissue is stimulated in the proximity of the recording site.

## 2 Materials and methods

### 2.1 Numerical simulations

Numerical calculations of optical scattering properties were performed by using finite-element method (FEM) numerical modeling in Comsol Multiphysics to simulate the propagation of light with a wavelength of 650 nm through the waveguide. The model cross-sections used were up to 4.5 × 4.6 µm; all the other geometrical parameters are scaled down proportionally to these values.

### 2.2 Preform fabrication

The preform for the thermal fiber drawing was 100× 18 × 12 mm prism with four holes along its length, produced by machining a PC cylinder with a starting diameter of 25 mm and a length of 100 mm. The diameter of the channels was 2.7 and 2 mm for the two side and central holes, respectively, to compensate for the partial collapsing of the side holes observed during previous thermal drawings. The preform was annealed at 50 °C in a vacuum dryer for 72 hours after machining.

### 2.3 Thermal fiber drawing

The preform was drawn by a standard thermal drawing process in an in-house facility. The furnace temperature was slowly ramped up to 212 °C, and an internal pressure of 6 mbar was set to avoid excessive collapsing of the channels in the waveguide during drawing. During drawing, the waveguide’s cross-sectional dimensions were controlled by tuning the capstan and feeder speed. The waveguide was subsequently cut into pieces of varying lengths (1.5 to 5 cm) for developing the devices.

### 2.4 Micromachining and connectorization

Four holes were micromachined on the top side of each device, in correspondence with the hollow microchannels, using a laser micromachining tool (microSTRUCT Vario). The tool was equipped with a 50 W picosecond laser (pulse duration = 12 ps) and a central wavelength converted from 1064 nm to 355 nm with a third-harmonic generation (THG) crystal. The power of the laser was decreased by setting a thin-film polarizer (TFP) power to 50% to better control the machined hole depth by changing the repetition rate of the laser. Following this fabrication step, tungsten microwires (longer than the waveguide pieces by several centimeters) of 50 µm in diameter were then manually inserted inside the four channels of the devices (Fig. 2e). On one end of the waveguide, the microwires were cut off and the channels were subsequently sealed and electrically insulated with cyanoacrylate adhesive. On the other end, the exposed microwires were connectorized with a 3D-printed adapter.

### 2.5 Optical characterization of the scattered light

The scattering was evaluated by coupling light from a supercontinuum source (SuperK Extreme) into the waveguide and then using a micrometric translational stage to scan the facet of a silica optical fiber (550 µm core diameter, 0.22 NA) along the top side of the device, with steps of 500 µm. At each position, the light collected by the fiber was measured using a silicon photodiode (Thorlabs S120 C).

### 2.6 Impedance spectroscopy

A chemical impedance analyzer (Hioki IM3590) was run in a three-electrode configuration to evaluate the electrodes’ impedance in 0.01 M phosphate-buffered saline (PBS) solution, in which the tungsten counter-electrode, the Ag/AgCl reference electrode, and a 3 cm-long optrode were immersed. For each measurement, a single electrode was clamped close to the electrical circuit and its impedance was measured in the 1-10000 Hz range.

### 2.7 Surgery

Electrophysiological recordings were performed in the spinal cord of Long-Evans wildtype adult rats. During the surgical operation, six vertebrae (11th thoracic through 2nd lumbar vertebrae) were exposed; the two outermost vertebrae were clamped, and the upper part of the bones was drilled in the four innermost vertebrae to expose the spinal cord. A 1.5 cm long device (in which the micromachined holes were separated by 2 mm) was connectorized with a pre-amplifier and an interface for electrophysiology (Intan Technologies RHD USB board) and a silica patch cable connected to a red LED (*λ* = 650 nm). The probe was then mounted on a motorized stereotaxic frame and lowered on the spinal cord. The electrophysiology setup was connected to a computer to display real-time data and record neural activity. All surgical procedures have been approved by the Animal Experiments Inspectorate under the Danish Ministry of Food, Agriculture, and Fisheries, and all procedures adhere to the European guidelines for the care and use of laboratory animals, EU directive 2010/63/EU.

### 2.8 Data analysis

All data processing, analysis, and plotting were performed in the Matlab suite.

## 3 Results and discussion

### 3.1 Numerical analysis of light propagation

Prior to the fabrication of the devices, their optical scattering properties were investigated by FEM. Due to computational constraints induced by a fine mesh and a large model geometry, the model was implemented by creating a down-scaled geometrical model with a reduced model cross-section of up to 4.5 × 4.6 µm. Part of the simulations with increasing model size showed that the smaller models overestimate the confinement losses of the waveguide; however, these models also supported a considerably reduced number of modes compared to the full-sized waveguide [21], which are the main contributors to the optical scattering induced by the micromachined holes and by the metal electrode interfaces. Furthermore, it is worth considering that our model does not account for additional scattering sources, such as surface roughness, additional air/metal interfaces in the four channels, or material impurities in the waveguide. Therefore, the scattering is expected to be significantly underestimated in the simulated results. Figure 1 reports the normalized scattered field distribution on a cross-section plane of the waveguide intersecting a hole for several horizontally-polarized modes. While the scattered power of the fundamental mode *n*_eff_ *≈* 1.59 is mainly oriented toward the bottom (Fig. 1a), as the mode order increases, the preferential scattering direction moves toward the sides of the waveguide. Nevertheless, the field distribution of Figure 1f shows that the features of the mode with *n*_eff_ *≈* 1.35 are small enough to concentrate in the proximity of the hole, which is confirmed by the increased scattering in the upward direction observed in the polar plot. The higher-order modes propagating in the full-sized waveguide are thus expected to further enhance the scattering on the upper surface of the waveguide, where the stimulation and the recording are projected to occur. Incidentally, other numerical simulations showed that as the size of the waveguide increases, the optical scattering of orthogonal polarizations converges. For small geometries (core width smaller than 2.5 µm), sideward scattering is slightly stronger in vertically-polarized modes, while upward and downward scattering is stronger in horizontally-polarized modes. However, in larger-sized simulations this difference gradually decreases; therefore, the polarization direction of the modes is not expected to produce any significant difference. These numerical simulations confirm the promising optical properties of the waveguide with rectangular cross-section for its intended use in multi-site optogenetic neurostimulation.

**Fig. 1.**
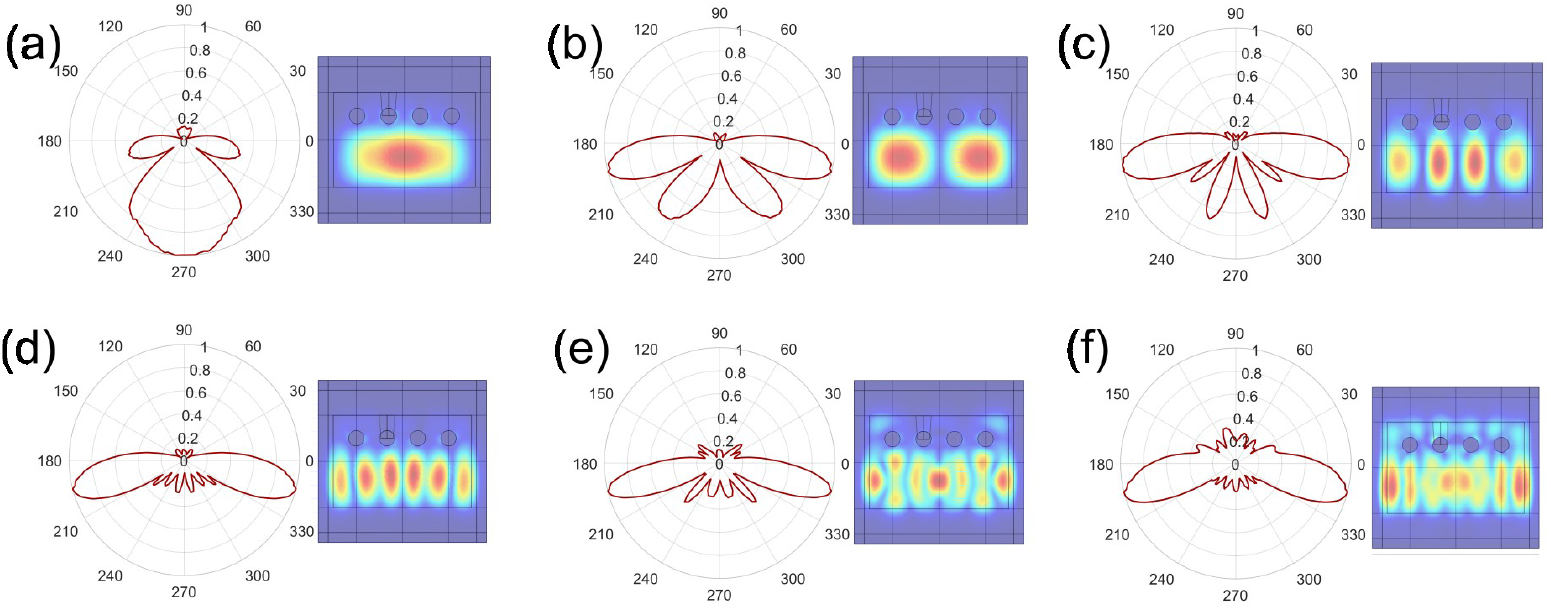
Normalized polar plots of the simulated electric field outside the waveguide, in the proximity of a hole, shown with the respective mode distributions on the side. The effective refractive indices around which the modes are searched are (a) 1.59, (b) 1.55, (c) 1.50, (d) 1.45, (e) 1.40, and (f) 1.35.

**Fig. 2.**
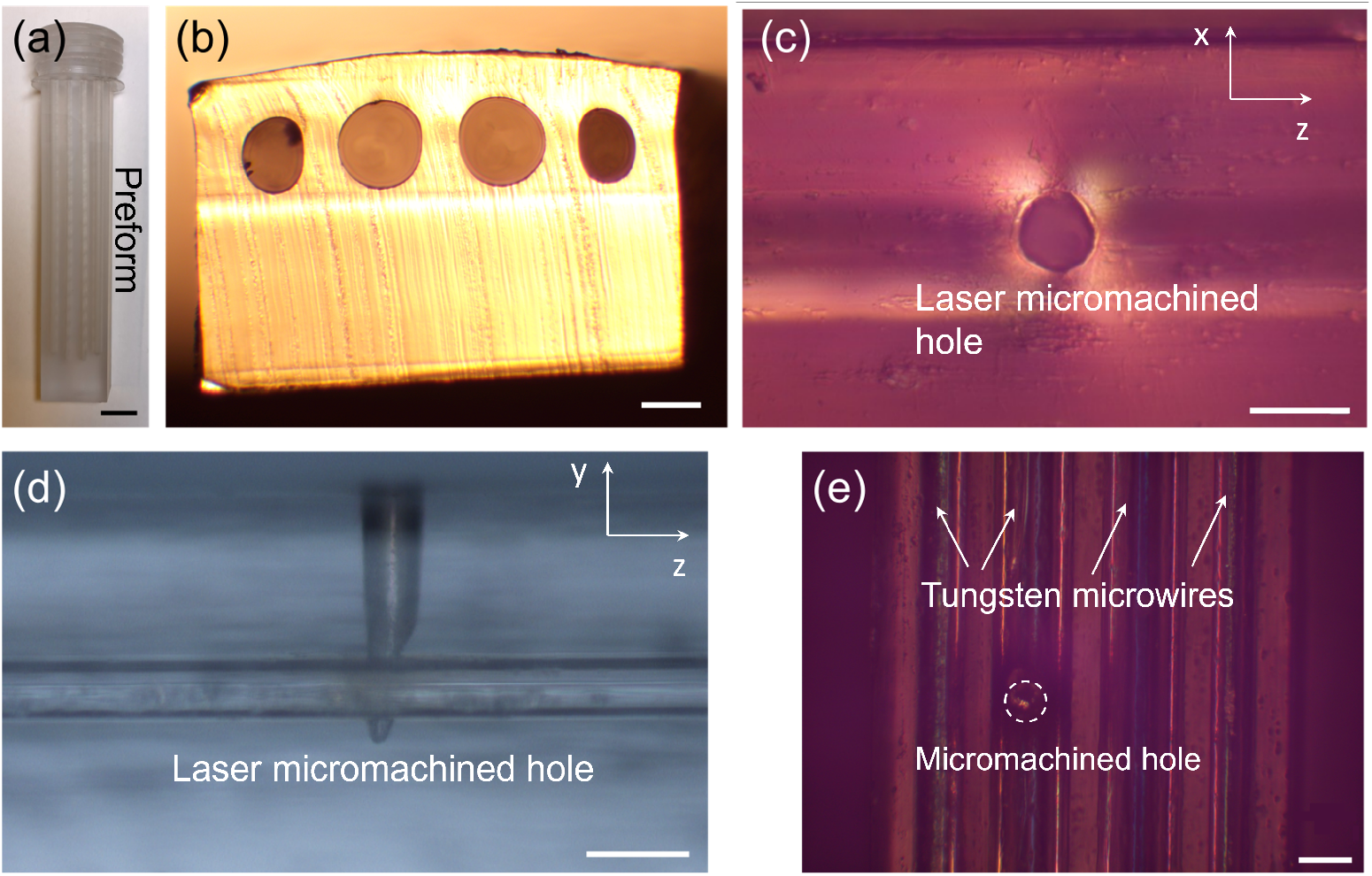
Fabrication of the PC-W optrode. (a) Photograph of the machined polycarbonate preform before thermal drawing; scalebar = 1 cm. (b) Optical microscope picture of the waveguide crosssection; scalebar = 100 µm. (c) Top-view picture of a hole from an optical microscope equipped with an analyzer, showing the cross-polarized scattered light by the edge of the hole; scalebar = 50 µm. (d) Side-view optical picture of a hole, exposing the underneath microstructured channel of the waveguide; scalebar = 100 µm. (e) Top-view optical picture of the waveguide after inserting the electrodes; scalebar = 100 µm.

**Fig. 3.**
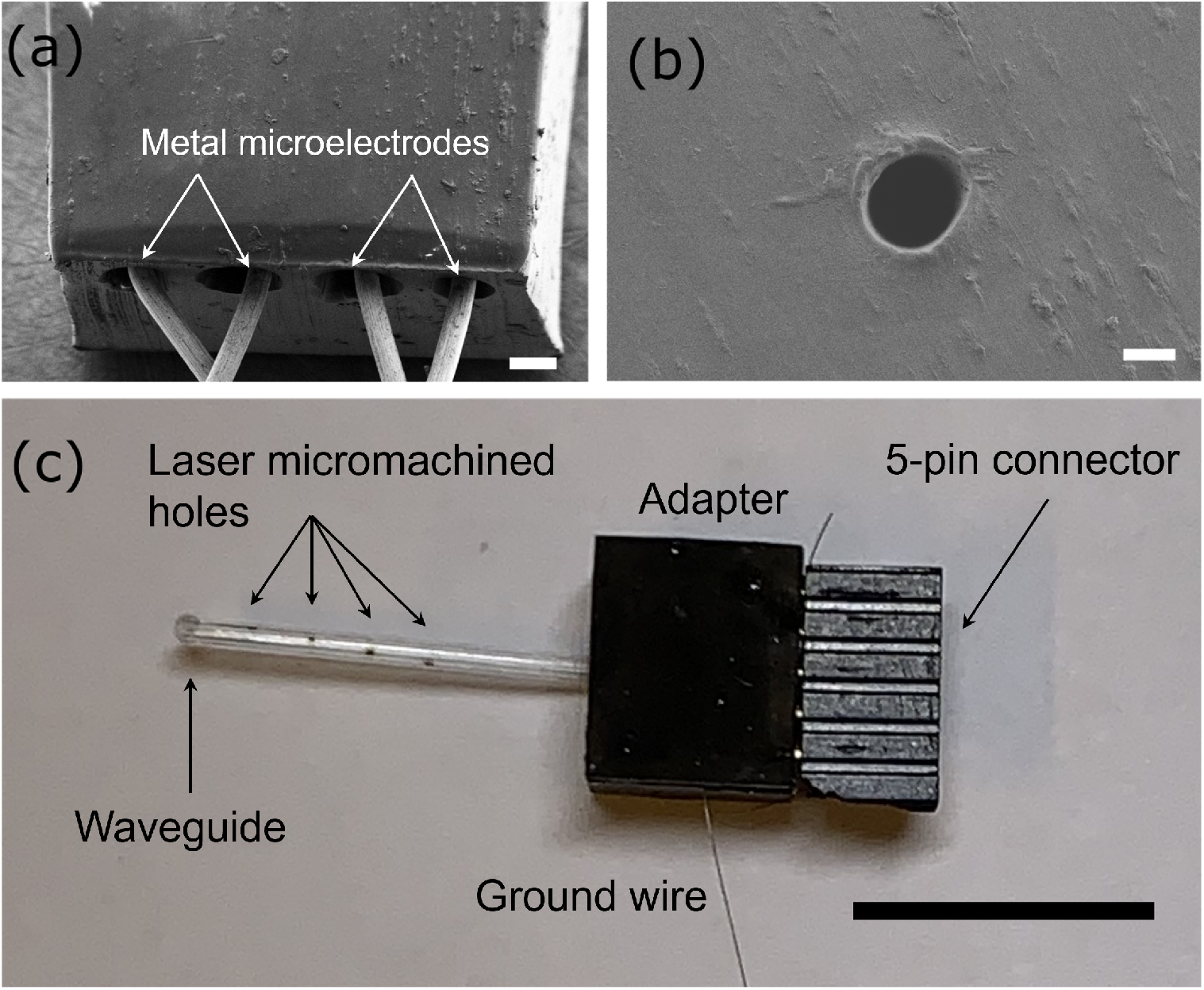
Image of the optrode, with (a) a scanning electron microscopy (SEM) image showing the end face of the waveguide (scalebar = 100 µm); (b) an SEM image of a micromachined hole on the surface of the waveguide (scalebar = 20 µm); (c) a full-scale photograph of the connectorized probe (scalebar = 1 cm).

### 3.2 Device fabrication and characterization

As a starting step in the fabrication of the proposed rectangular multi-site bidirectional spinal cord interfaces, a macroscopic template of the waveguide (preform) with hollow channels (Fig. 2a) was scaled down to sub-millimeter size by a standard thermal drawing process. The resulting waveguide maintained the desired shape to a high degree, with only a minor excess of collapsing in the sidemost holes attributed to the non-centrosymmetric geometry (Fig. 2b). Following the drawing, waveguide pieces of lengths ranging from 15 to 50 mm were cut and micromachined with holes to achieve contact between the tissue and the hollow channels (Fig. 2c). The parameters of the laser micromachining tools (e.g., laser power, jump delay, on/off delay, and horizontal/vertical alignment) were optimized to ensure a hole depth sufficient to reach the channels and an accurate repeatability (Fig. 2d). The final profile of the holes measured 85 µm in diameter and 175 µm in depth, with less than 4% variation between holes.

Finally, 50 µm diameter tungsten microwires were inserted along the whole length of the hollow channels to act as electrodes for electrophysiological recording (Fig. 2e), and the channels were then sealed at one end. Two scanning electron microscopy (SEM) images of the integrated metal microwires and the top hole are shown in 3a and b, respectively. The devices were completed by back-end connectorization using an inhouse-developed 3D printed casing with a 5-pin connector port for electrophysiology (the additional pin is connected to a fifth tungsten wire for electrical grounding) and a circular port for coupling light inside the waveguide through an optical ferrule. A photograph of the complete assembled neural interface is shown in 3c. Despite the additional bending stiffness introduced in the PC waveguide by the insertion of metallic microwires, the completed devices maintained a high flexibility, and could easily be bent at 1 cm bending radius (Fig. 4a). Furthermore, according to previous simulations from our group, we expect this additional stiffness to be mainly affecting the devices in the direction parallel to the longer side of the cross-section and to have only a minor effect on the bending required to conform the device to the curvature of the spine [22].

**Fig. 4.**
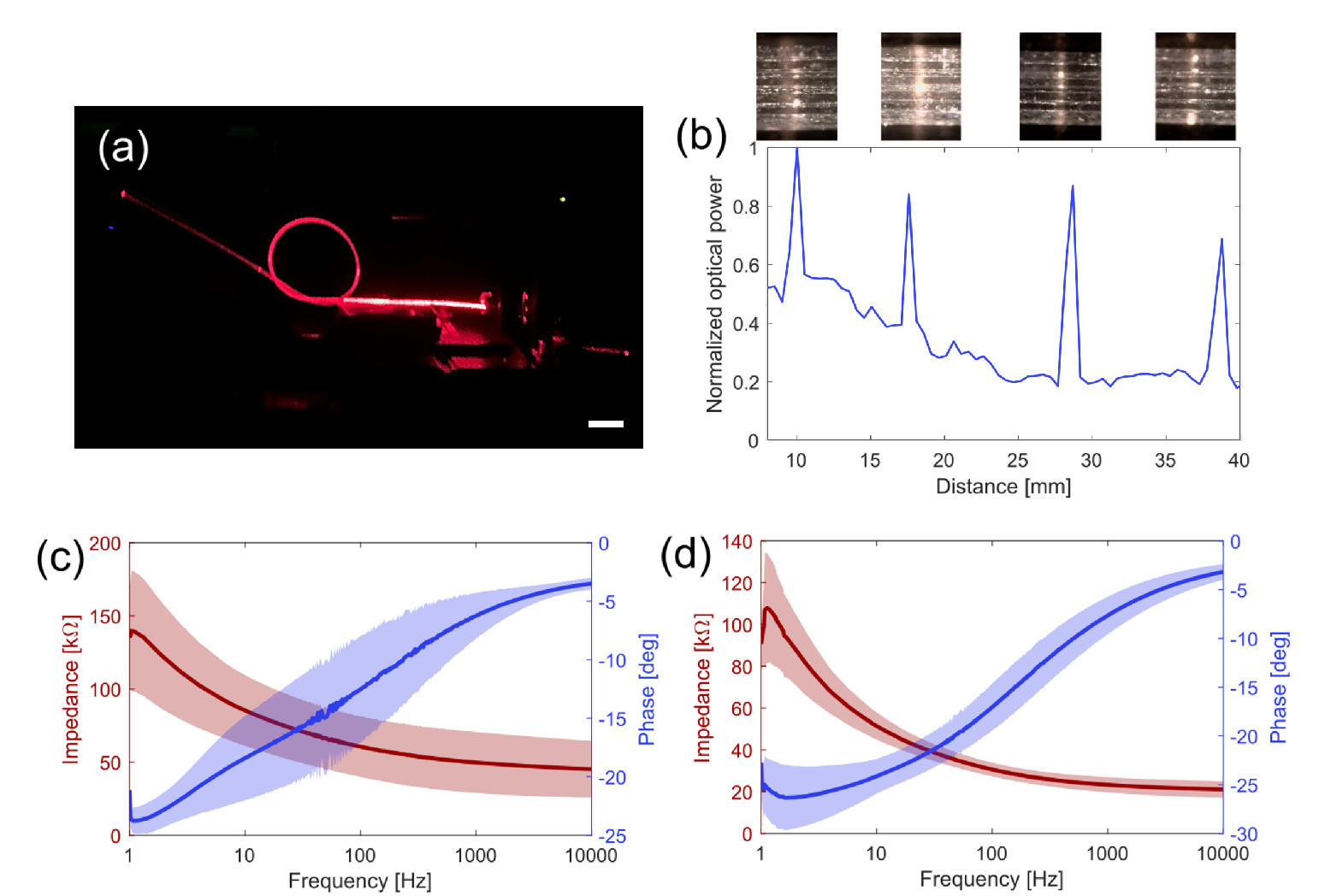
Optical and electrical characterization of the waveguide. (a) Flexibility of the waveguide, coupled to red laser light; scalebar = 1 cm. (b) Space-resolved optical scattering on the top surface of the waveguide after the insertion of tungsten microwires. (c) Electrochemical impedance spectroscopy measurements after 20 min; the shaded areas show one standard deviation. (d) Electrochemical impedance spectroscopy measurements after 24 h; the shaded areas show one standard deviation.

The scattering properties of the holes micromachined on the device were characterized by space-resolved power measurements of the light collected by a silica optical fiber scanned along the top side of a 5 cm-long waveguide in air, after the insertion of tungsten microwires (Fig. 4b). The four peaks, each corresponding to a micromachined hole (as shown in the insets, displaying optical microscope pictures of scattered white light from the top of the probe), show a clear contrast with the background scattered light, demonstrating the possibility of local tissue stimulation by adjusting the input power so that only the fluence at the scattering sites’ locations will reach the activation threshold for the employed photoreceptors. The impedance of the four individual recording electrodes was evaluated using electrochemical impedance spectroscopy (EIS). The different electrodes (corresponding to the different holes along the device) gave similar impedance values; hence, the measured spectra were averaged and shown with the error for two sets of measurements: one shortly after the submersion of the device in the PBS solution (Fig. 4c), and one after 24 hours (Fig. 4d), when the redox species had stabilized on the electrode surface, thereby giving a considerably lower error between measurements (decreased on average by 42% for the impedance magnitude and by 25% for the phase). Nevertheless, in both cases, the recorded magnitude of the impedance is comparable to state-of-the-art recording probes featuring metallic electrodes with small dimensions [23–25].

### 3.3 *In vivo* spinal cord electrophysiological recordings

To validate the capability of our optrodes to perform spinal cord recordings, we conducted an *in vivo* electrophysiology experiment on a rat in acute settings. This was performed by using a 1.5 cm long device in the exposed spinal cord of the rat, between the 11^th^ toracic and 2^nd^ lumbar vertebrae (Fig. 5a,b). At this stage, a viral vector for optogenetics had not been injected into the rodent due to time constraints in the project; nevertheless, red light was coupled into the device to demonstrate the visibility of its four scattering points even by eye (inset of Fig. 5c). Due to the design of the device, one could expect strong photoelectric artifacts to arise during optogenetic stimulation when light scattered by the holes hits the conducting microwires. However, several strategies can be applied to mitigate this issue, such as the use of red-shifted opsins to reduce photon energy, the use of high-frequency pulse trains followed by low-pass filtering in local field potential (LFP) recording, or the exploitment of the artifacts’ temporal consistency to remove them during signal post-processing [26].

**Fig. 5.**
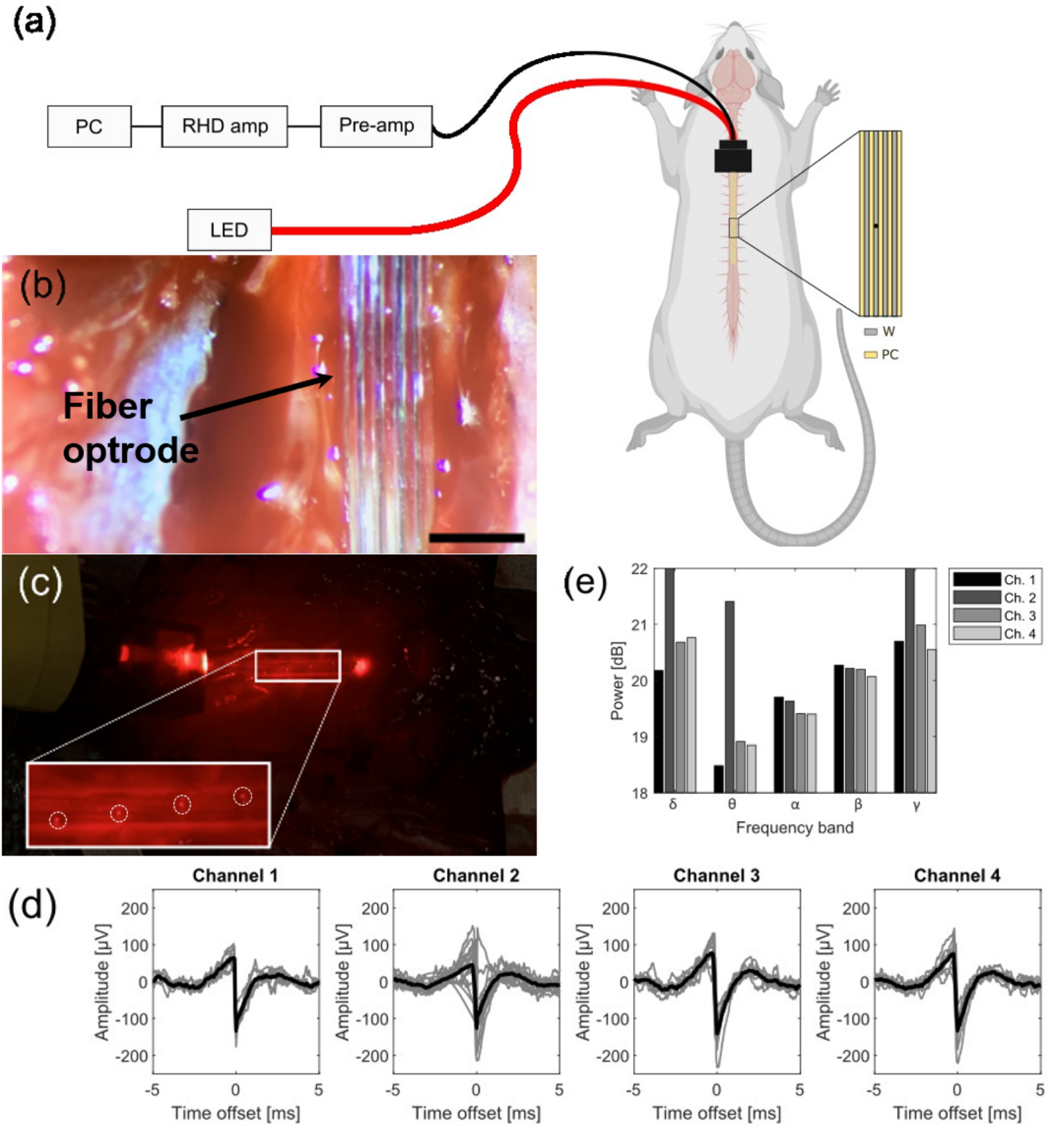
*In vivo* acute electrophysiology on an anesthetized rat. (a) Schematic of the setup for electrical recording and optical stimulation on the rat, where the spinal cord is emphasized. (b) Optical microscope picture of a portion of the probe on the surface of the spinal cord. (c) Picture of the probe mounted on a stereotaxic, showing scattered light near the micromachined holes; the inset shows a magnified view of the four stimulation and recording sites. (d) Recorded action potentials in the four channels (gray) and average signal of a spike (black). (e) Neural oscillations divided into spectral bands for local field potentials.

A series of nine traces at different positions in the spinal cord, each trace consisting of a 3-minute recording, was collected. The electrophysiology traces were then rectified with a high-pass filter at 200 Hz and then scanned for action potentials between*™* 300 and 300 µV in amplitude. The observed signals (Fig. 5d) match the expected spike shape for action potential in the spinal cord [27], with some exceptions in Channel 2, which exhibits two spurious spikes. A second analysis focused on the lower-frequency band, i.e. LFPs. By calculating the power spectral density and integrating the spectral power into the frequency bands for neural oscillations, we observed high activity for delta waves (0-3 Hz) and low activity for theta waves (4-7 Hz) compared to alpha (8-13 Hz), beta (13-30 Hz), and gamma (30-150 Hz) waves in all channels except the second, which, as previously observed, had recorded the highest noise among the four channels (Fig. 5e). Strong delta brainwaves and suppression of theta activity have been associated with the loss of consciousness induced by anaesthetics [28]; hence, the results are in agreement with the state of the rodent during the acute experiment.

## 4 Conclusions

In conclusion, we have developed the first, to the best of our knowledge, flexible optrode design for multi-site stimulation and recording specifically targeting the spinal cord. The novel design for the optrode is expected to open up new possibilities for enabling distributed optical stimulation and high-density, space-resolved electrical recordings. The results from the numerical modeling have shown enhanced upward scattering for higher-order modes, which could be tuned further via numerical optimization of the micromachined geometry. The fabrication process, based on the thermal drawing of flexible polymers and the insertion of tungsten microwires, enables cost-effective and scalable production of optrodes, which can potentially be further scaled down in size by optimizing the thermal drawing and decreasing the diameter of the microwires, to improve the flexibility of the probe, the response of the electrodes and the ease of implanting the probe. To validate the optical stimulation properties of the optrode, future work should be focused on testing the probe with a photostimulation method, i.e. either optogenetics [6, 8] or infrared neural stimulation [25].

## Acknowledgments

The authors would like to acknowledge financial support from Lundbeckfonden (Multi-BRAIN project R276-2018-869 and R380-2021-1171). The authors would also like to acknowledge support from DTU Electro Internal Projects and DTU Nanolab for providing access to the equipment.

## Disclosures

The authors declare no conflicts of interest.

## Data availability

Data underlying the results presented in this paper are not publicly available at this time but may be obtained from the authors upon reasonable request.

## Notes

### Competing Interest Statement

The authors have declared no competing interest.

